# Schild analysis of the interaction between parthenolide and cocaine suggests an allosteric relationship for their effects on planarian motility

**DOI:** 10.1101/2022.01.02.474598

**Authors:** Jyothi Kakuturu, Mary O’Brien, Oné R. Pagán

**Affiliations:** Department of Biology, West Chester University

**Keywords:** planaria, cocaine, parthenolide, motility, Schild analysis, allosteric, orthosteric

## Abstract

The freshwater planarian is an emerging animal model in neuropharmacology due to its close parallels with the nervous system of vertebrates. Cocaine is a naturally occurring compound and an abused drug, as well as the ‘founding member’ of the local anesthetic family. Sesquiterpene lactones of the parthenolide class are naturally occurring products that act as behavioral and physiological antagonists of cocaine in planarians and rats, respectively. One of the best-characterized planarian behaviors induced by various compounds is the change in locomotor velocity. Previous work from our laboratory showed that both parthenolide and cocaine reduced planarian motility and that parthenolide reversed the cocaine-induced motility decrease at concentrations where parthenolide does not affect the movement of the worms. However, the exact mechanism of the cocaine/parthenolide antagonism is unknown. Here, we report the results of a Schild analysis to explore the parthenolide/cocaine relationship in the planarian Girardia tigrina. The Schild slopes of the Δ parthenolide ± cocaine and Δ cocaine ± parthenolide were -0.33 and -0.43, respectively. These slopes were not statistically different from each other, and both were statistically different from -1, suggesting an allosteric relationship between these two compounds. To the best of our knowledge, this is the first study aimed at studying the mechanism of action of the antagonism between cocaine and parthenolide.

## 1. Introduction

Freshwater planarians have a distinguished history in regenerative and developmental biology research; the study of these organisms in this context has been ongoing for more than two centuries and is still a source of profound insights in these areas, particularly since the availability of molecular biology, genomics, and bioinformatics tools and techniques [1-7]. More recently, planarians have emerged as a powerful model to study regenerative processes at the molecular level, specifically in nervous tissue (reviewed in [8-12]). Planarians are also an emerging pharmacological model system for three main reasons: (1) the varied behavioral responses that these organisms exhibit upon exposure to exogenous substances, responses closely reminiscent of vertebrate responses to drugs, (2) the similarities of their nervous system to the nervous system of vertebrates in terms of cellular biology and morphology, and (3) the presence of vertebrate-like neurotransmitter systems in these organisms [13-17]. An especially active area of pharmacological research using planarians is the in vivo study of the effect of abused drugs, including cocaine, amphetamines, ethanol, and nicotine, among others, to explore their effects in terms of acute exposure (toxicity) or chronic administration, meaning sub-lethal concentrations that may lead to phenomena like desensitization, tolerance, and withdrawal-like behaviors [18-29]. Thus, planarians are uniquely positioned to integrate the ostensibly separate disciplines of regeneration, neuroscience, and pharmacology [30,31].

Cocaine (Figure 1) is a plant-derived compound (mainly from plants of the Erythroxylum genus [32,33]) and is the parent compound of the local anesthetic superfamily, which primarily targets voltage-gated sodium channels on the surface of nerve cells [34]. Besides its local anesthetic properties, cocaine’s main behavioral effect in vertebrates is through its interaction with neurotransmitter transporters, particularly the dopamine transporter [35]. The naturally occurring sesquiterpene lactone parthenolide (Figure 1) is mainly isolated from the feverfew plant (Tanacetum parthenium, [36,37]), a plant used in traditional medicine in various cultures. Parthenolide and other sesquiterpenoid lactones are currently in clinical trials for their antitumoral and anti-inflammatory properties [38-41], among other interesting biochemical effects. However, parthenolide and related compounds also induce contact dermatitis and genotoxic effects [42-44].

**Figure 1.**
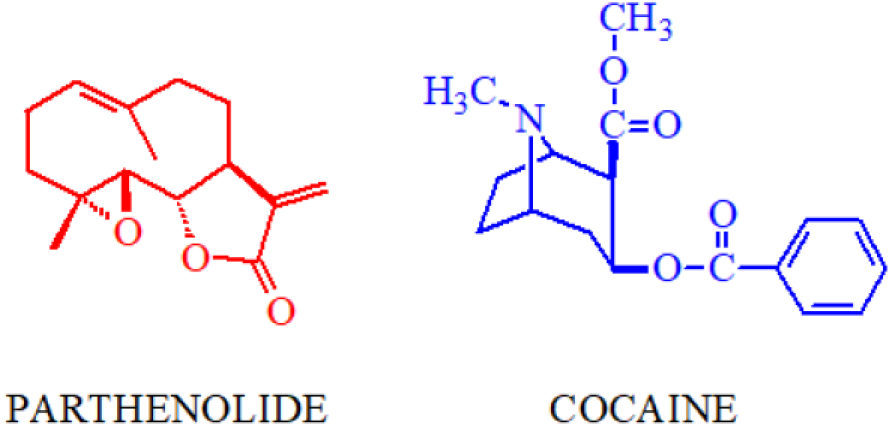
Compounds studied in this work.

The logical thread that led us to the study of the interaction of cocaine and the sesquiterpene lactones of the parthenolide family and the initial use of planarians for these purposes is reviewed in detail elsewhere [4,45]. Our research established that parthenolide is a behavioral antagonist of cocaine in the planarians system, capable of counteracting the effects of acute cocaine exposure, which mimic toxicity effects [46], as well as preventing the induction of withdrawal-like behaviors by cocaine, which mimic chronic administration [47]. We also determined the minimal structural determinants of the parthenolide-like molecules capable of antagonizing the motility decrease induced by cocaine [48]. Additionally, we established that parthenolide is a specific cocaine antagonist for another acute administration effect, the induction of seizure-like movements [49] through experiments in which parthenolide prevented the expression of seizure-like behavior by cocaine but did not prevent the expression of similar behavior induced by nicotine and amphetamines, among other drugs [50]. Furthermore, the research group of a colleague and collaborator, Dr. Carlos Jiménez-Rivera (University of Puerto Rico) found that parthenolide antagonized the inhibitory effect of cocaine on neuronal firing rate in the ventral tegmental area of rats, opening the door to the potential of parthenolide as a cocaine antagonist in vertebrates [51] validating the usefulness and vertebrate relevance of the planarian model.

Although the molecular targets of cocaine in mammals are well-understood, little is known about the mechanism of action and molecular nature of the cocaine target (s) in planarians. In the present work, we explore the relationship between parthenolide and cocaine in terms of their effects on planarian motility using a Schild analysis to obtain information about the mechanism of this interaction.

## 2. Materials and Methods

Brown planarians (Girardia tigrina) were purchased from Ward’s (Rochester, NY). Parthenolide and cocaine were purchased from Tocris (Minneapolis, MN) and Sigma-Aldrich (St. Louis, MO), respectively. General laboratory materials, chemicals, and supplies were purchased from Fisher Scientific (Suwanee, GA). All graphs and statistical analyses were done with the Prism software package (GraphPad Inc., San Diego, CA). Upon arrival at our laboratory, the flatworms were transferred to artificial pond water (APW: 6 mM NaCl, 0.1 mM NaHCO3, 0.6 mM CaCl2) and were allowed to acclimate to laboratory conditions for at least 24 hours before using them for experiments. The worms were used within three weeks of arrival. The APW was changed at least once every day except during weekends and immediately before any experiments. All experiments were performed at room temperature in APW containing 0.1 % dimethylsulfoxide (DMSO) as a solubility-enhancing agent for parthenolide; this concentration of DMSO does not display any apparent behavioral or toxic effects in planarians [52].

We adapted a published protocol to measure planarian motility [53], as modified in [24,25,46,52,54]. Briefly, using a small paintbrush or plastic pipette, a planarian (1.0-1.5 cm long) was placed in a transparent, APW-rinsed, 6 cm polystyrene Petri dish. The dish was placed over a one cm2 grid, and five milliliters of APW/0.1 % DMSO (control) or 5 mL of the experimental solutions in 0.1 % DMSO were added to the dish. The motility of the worm was measured by counting the squares crossed or recrossed per minute for 5 minutes. Each worm was used only once. The data was then plotted as cumulative crosses vs. time and fit using simple linear regression to obtain its slope. In experiments where the worms were exposed to various concentrations of the experimental compounds, the slopes obtained from the linear regression were normalized to control slopes and graphed as the fraction of control vs. the experimental compound concentration [24,25,46,52,54]. To analyze these concentration-response curves, we fit the data to an empirical Hill equation (Equation 1).

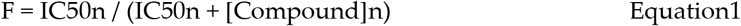

F is the fraction of control, [Compound] is the experimental compound concentration in μM, the IC50 is the compound concentration that decreased planarian motility by 50 %, and n is the Hill coefficient. The Hill coefficient is a parameter that can be used to estimate the kind and the degree of cooperativity in multimeric proteins. Although a useful parameter for the initial exploration of cooperativity, its direct link to the actual physical reality of a system is unclear [55,56]. In general, the usefulness of empirical models lies in the fact that they may provide helpful mechanistic insights and information about the pharmacology of a compound when little or no information exists about either the ligand or its putative target [57-59].

We performed two sets of experiments to observe the effect of parthenolide or cocaine on the IC50 of the other to decrease planarian motility: A series of concentration-response curves of cocaine in the absence or the presence of single concentrations of parthenolide (Δ cocaine ± parthenolide), and a series of concentration-response curves of parthenolide in the absence or the presence of single concentrations of cocaine (Δ parthenolide ± cocaine). In these experiments, the compound used in the concentration-response curve (the Δ compound) was defined as ‘Compound A,’ and the compound added at a single concentration was defined as ‘Compound B.’ We used concentrations of compound B that did not slow down the worms [50]. The shift in the IC50 for compound A induced by the single concentration of compound B was subjected to a Schild analysis by fitting the data to a linear equation (Equation 2).

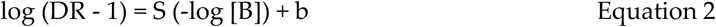

The DR is the dose ratio (the IC50 value obtained from a concentration-response curve of compound A in the presence of a single concentration of compound B over the IC50 value of compound A in the absence of compound B). S is the Schild Slope, and b is the y-intercept. A Schild Slope statistically equal to unity suggests an orthosteric relationship between the tested compounds; conversely, a Schild Slope statistically different from unity suggests an allosteric relationship between the tested compounds [60-65].

## 3. Results

Figure 2 shows the concentration-response curves of cocaine or parthenolide for planarian motility. The IC50 values (421 and 93 μM for cocaine or parthenolide, respectively) are consistent with previous reports from our laboratory [50].

**Figure 2.**
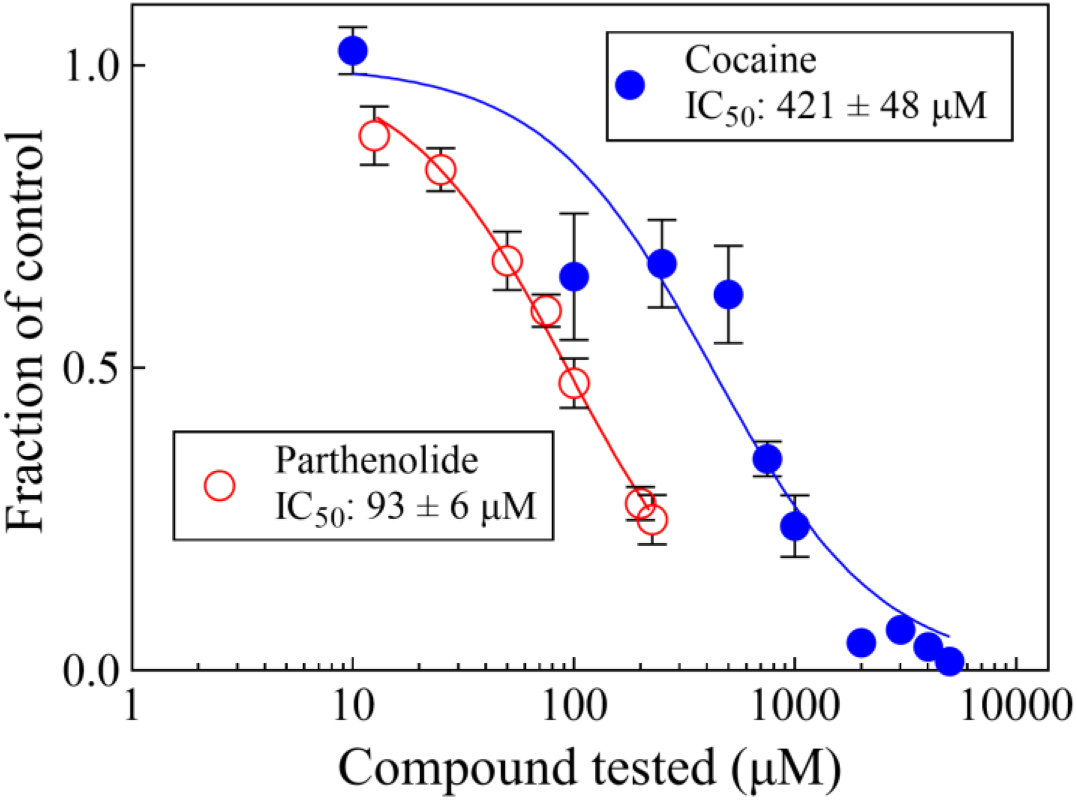
Parthenolide and cocaine decrease planarian motility in a concentration-dependent manner. Each data point represents the average of the responses of 6 worms. The lines were generated by fitting the data to Equation 1. The error bars represent the standard error of the mean. Symbols without visible error bars are bigger than the size of the SEM value.

Figure 3 shows representative curve pairs of Δ parthenolide ± cocaine or Δ cocaine ± parthenolide, showing the apparent IC50 shift of the experimental compounds, as indicated. Pairs of curves for each of the conditions were obtained, and their parameters from the fit to either Equation 1 or Equation 2 are shown in Table 1.

**Table 1.**
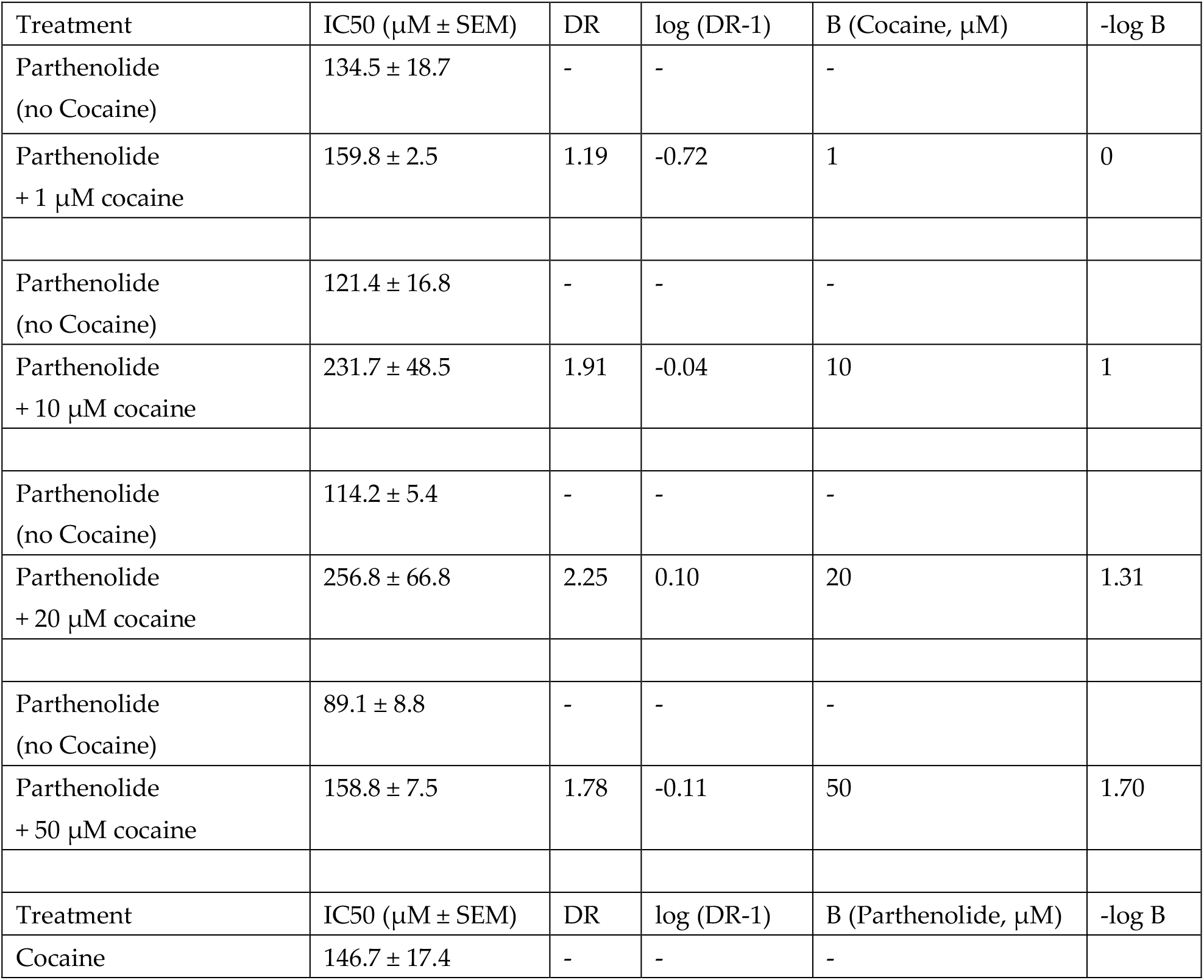
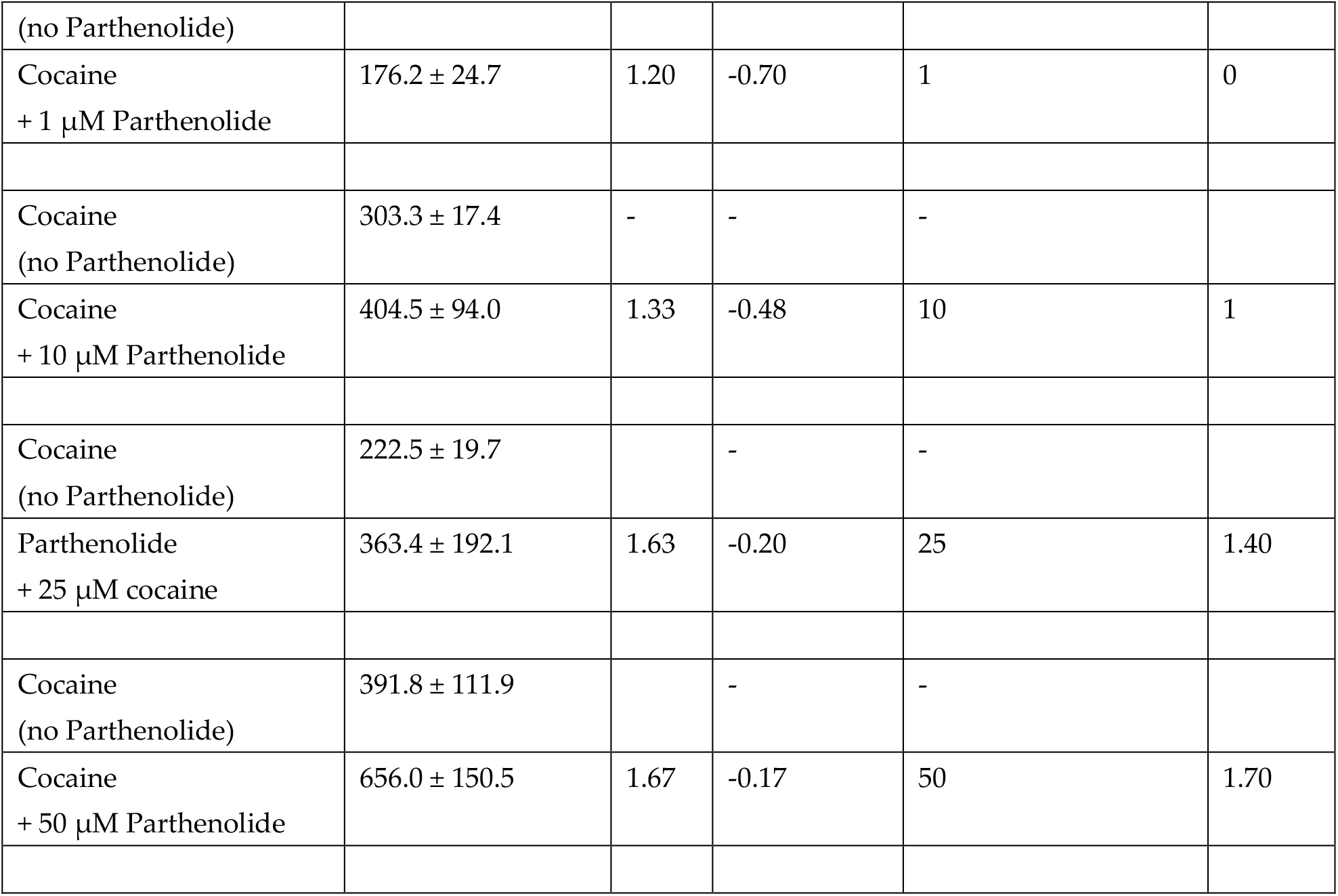
Hill and Schild parameters for the data fit to Equations 1 and 2 (see text).

**Figure 3.**
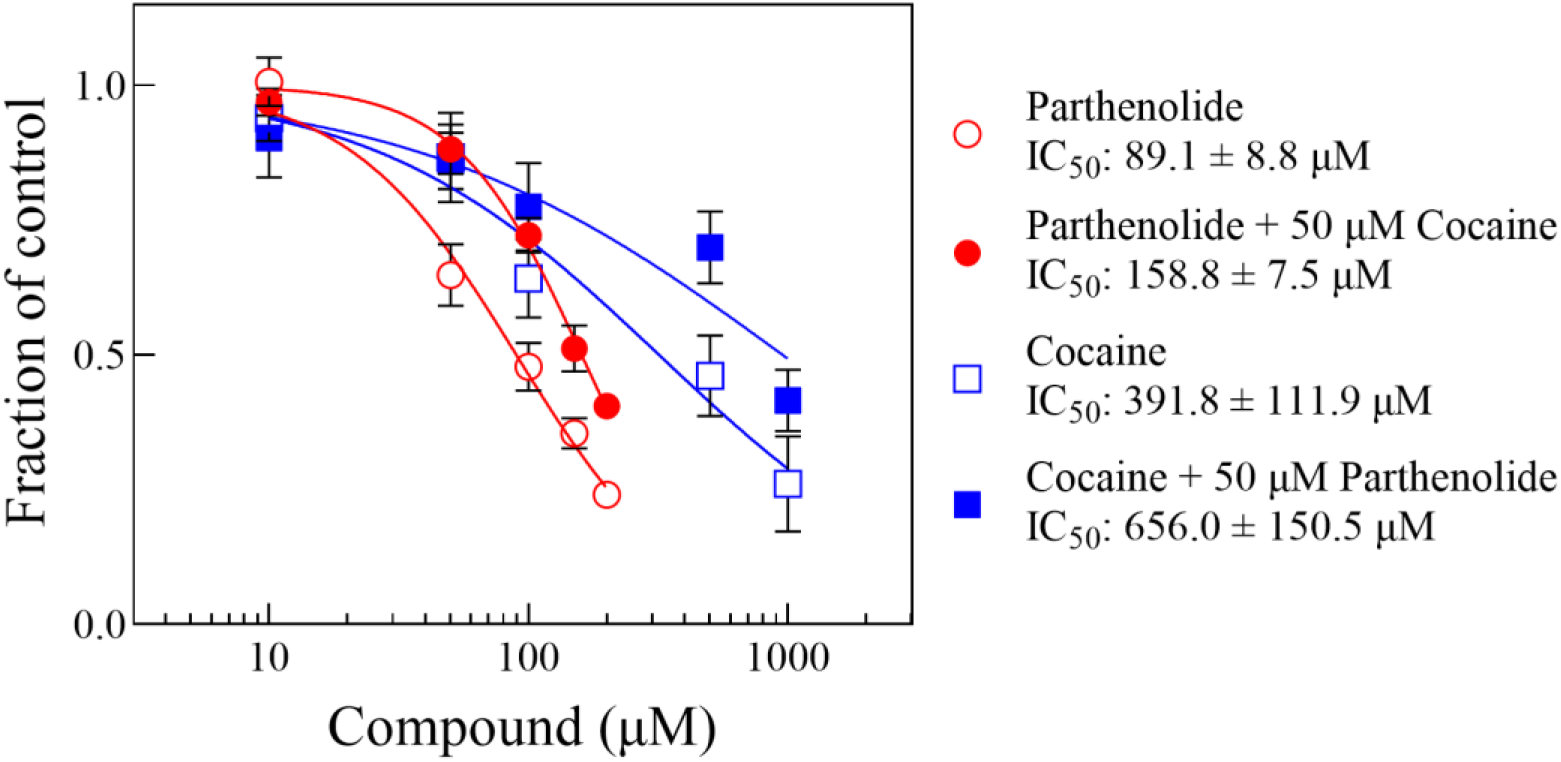
Representative concentration-response curves of motility decrease by parthenolide or cocaine in the absence and the presence of a single concentration of the other, as indicated (see text and Table 1). Each data point represents the average of the responses of 4 worms. The lines were generated by fitting que data to Equation 1. The error bars represent the standard error of the mean. The Hill and Schild parameters for all pairs are listed in Table 1. Symbols without visible error bars are bigger than the size of the SEM value.

Figure 4 shows Schild plots of the two tests we performed. For the Δ parthenolide ± cocaine or of Δ cocaine ± parthenolide the Schild slopes were -0.33 ± 0.06 and -0.43 ± 0.18, respectively. Both slopes were statistically different from -1 (p = 0.002 and 0.0498, respectively, one sample t-test) and were not statistically different from each other (p = 0.633, unpaired t-test).

**Figure 4.**
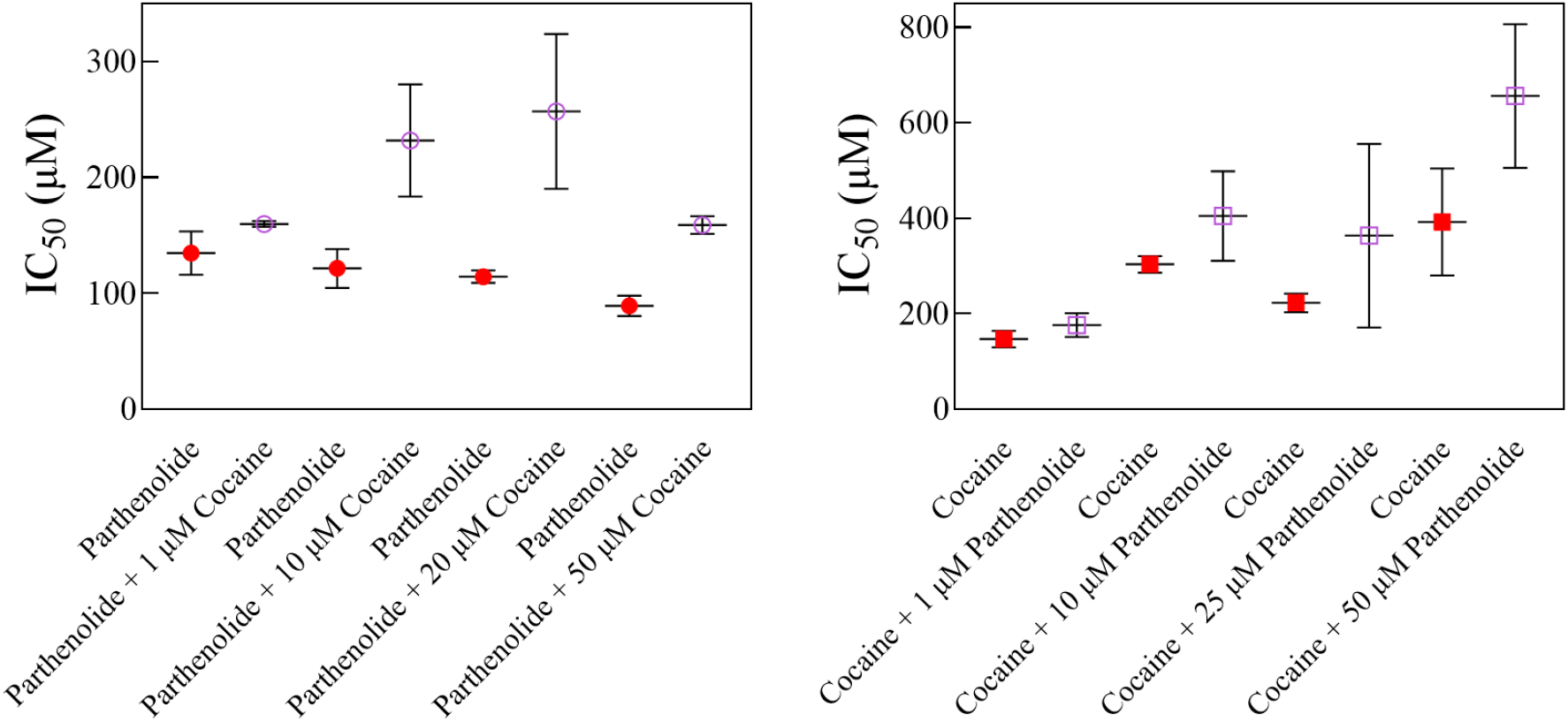
Parthenolide increases the apparent IC50 of cocaine and vice versa. The scatterplots of Δ parthenolide in the absence or in the presence of cocaine (at left) or Δ cocaine in the absence or in the presence of parthenolide (at right) show that for each treatment pair, the IC50 of the Δ compound increases with the presence of the second compound (see text). Error bars represent the standard error of the mean.

## Discussion

In this work, we present an initial analysis of the parthenolide and cocaine antagonism in the planarian model. The molecular targets of cocaine in vertebrates are well known (voltage-gated sodium channels and monoamine transporters), and its interaction with these targets accounts for virtually every observed effect of cocaine in biological systems. Planarians possess genes analogous to these proteins according to genomic data available in various databases [6,7,66,67]. All available evidence suggests that these proteins function in planaria as in vertebrates. In contrast, the nature of the molecular target (s) of parthenolide and related sesquiterpene lactones is less clear; in fact, parthenolide has been described as having a “…promiscuous bioactivity profile…” [40]. Most studies on the molecular effects of parthenolide and related compounds focus on their impact on IkB kinases, the nuclear factor kappa-B (NF-κB), and several classes of cytokines downstream, which may account for parthenolide’s antitumoral and anti-inflammatory properties [68,69]. Parthenolide, in particular, seems to decrease NF-κB activity at the transcriptional level and by direct inhibition of IKK-B kinases [39].

NF-κB proteins seem to be a common target for sesquiterpene lactones and cocaine. Cocaine increases NF-κB activity in H9C2 cells (striated cardiomyocytes from rats, [70]), and mice chronically exposed to cocaine display enhanced activity of NF-κB pathways [71]. Additionally, sodium diethyldithiocarbamate trihydrate (DTT), a NF-κB inhibitor, reverses the cocaine-induced expression genes related to axon guidance pathways in mice [71]. Paradoxically, long-term cocaine exposure induces NF-κB activity in mice hippocampus [72]. Interestingly, the viral knockdown of NF-κB within the rat nucleus accumbens induces a reduction in cue-induced cocaine seeking in males but not female rats [73], and chronic cocaine administration induces NF-κB-dependent transcription in mice [74]. There are analogs of NF-κB proteins in various planarian species [6,7,66,67]. Although an intriguing possibility, the apparent role of NF-κB proteins as a link between parthenolide and cocaine is speculative at the moment. More studies are needed to explore this possibility. That said, based on our results, even if NF-κB is the common target of parthenolide and cocaine in the planarian model, the most likely nature of this relationship would be a common pathway rather than direct binding sites for both experimental compounds on the protein.

The fact that cocaine and parthenolide affect the apparent effect of the other indicates a mutually exclusive binding relationship between these compounds for their motility decrease properties in planarians. This interpretation is consistent with previous studies from our laboratory [46] (Figures 3 and 4). However, the exact nature of this relationship is unknown. The most straightforward interpretation of our data would be that parthenolide and cocaine display an orthosteric interaction (a single type of binding site shared by both compounds and, alternatively, separate yet overlapping binding sites). On the other hand, the parthenolide/cocaine interaction could be allosteric, where each compound would have its individual, non-overlapping (yet related to the other’s) binding site. To distinguish between these two possibilities, we performed a Schild analysis [60-61] of the IC50 shift data from Table 1. Briefly, if the value of the Schild slope is statistically equal to unity, the data would suggest an orthosteric relationship between parthenolide and cocaine. On the other hand, if the slope is significantly different from unity, the data would suggest an allosteric relationship between the two compounds [59-65]. Our analysis found that Schild Slopes or parthenolide and cocaine were -0.33 ± 0.06 and -0.43 ± 0.18, respectively. Both slopes were statistically different from -1 (p = 0.002 and 0.0498, respectively) and were not statistically different from each other (p = 0.633), suggesting that the parthenolide/cocaine relationship in our model is allosteric in nature.

In planarians, cocaine induces other types of behaviors in addition to motility decrease. Some of these behaviors are expressed under acute exposure conditions, such as seizure-like hyperkinesia [26,27,31] (an effect that parthenolide also alleviates [50]), dark/light environmental preference (which is oftentimes interpreted as an anxiety-like response [75,76]), and behavioral and cross-sensitization [77]. Also, chronic exposure to cocaine induces conditioned place preference behaviors [78] and even withdrawal-like behaviors [28,29] (a type of behavior that parthenolide does not induce; [47]). Of these cocaine-induced behaviors, we have demonstrated that parthenolide and related analogs antagonize cocaine-induced motility decrease and seizure-like hyperkinesia [46,50], as well as withdrawal-like behaviors [47].

Microtubules are another possible protein candidate that may be a common target of parthenolide and cocaine. Parthenolide and costunolide interfere with microtubule physiology [79,82], and microtubules and microtubule-associated proteins are “non-canonical” targets of cocaine [83-85]. Interestingly, parthenolide seems to promote functional nervous tissue regeneration in mice [86]; this contrasts with results obtained from our laboratory that suggest that parthenolide slows down the rate of planarian brain regeneration (manuscript in preparation).

Additional studies that might contribute to the understanding of the parthenolide/cocaine relationship in planarians may include detailed mechanistic analyses of cocaine-induced seizure-like hyperkinesia, dark/light environmental preference, withdrawal-like behaviors, and habituation and desensitization. As discussed previously, cocaine induces such effects in planarians. These studies will help paint a clearer picture of the relationship between our experimental compounds, especially since it is likely that cocaine induces such effects by affecting different molecular targets. The study of these other cocaine-induced behaviors in planarians and their alleviation by sesquiterpene lactones of the parthenolide class will likely shed light on the mechanism of action of cocaine and perhaps other related abused drugs. Furthermore, an exciting possibility would be to conduct detailed structure-activity studies of cocaine alleviation by different sesquiterpene lactones of the parthenolide class. These studies may prove valuable for understanding this pharmacological problem. We have preliminary information in this respect. We established that a lactone moiety is necessary for an anti-cocaine effect [46,48] and that costunolide, a close structural analog of parthenolide, is about 1.6 times more potent for the alleviation of cocaine-induced motility decrease in planarians [46]. In contrast, santonin, another sesquiterpene lactone (Figure 6), does not alleviate the cocaine-induced motility decrease in planarians [46]. This information showcases the potential of structure-activity studies in the present context, as it is evident that relatively small structural differences significantly affect the potency of sesquiterpene lactones in this context, especially considering the close structural similarity between parthenolide and costunolide. The only difference between these two compounds is that a double bond substitutes the epoxy group in carbons 4 and 5 of parthenolide, while santonin displays additional structural differences compared to parthenolide and costunolide (Figure 6).

## 6. Conclusions

Our work indicates that parthenolide and cocaine are inhibitors of each other in the planarian system, consistent with previous reports from our laboratory [46]. Further, our results indicate an allosteric relationship between these compounds based on a Schild analysis discussed above. However, in the experiments of Δ cocaine ± parthenolide, the Schild slope was on the edge of statistical significance (p = 0.0498, Figure 5). Further experiments may include additional concentration-response curves of both compounds in the absence and the presence of the other, which should fine-tune our results and provide more statistical power to our experiments. Any additional information that we obtain from this approach, as well as studies on other cocaine-induced behaviors in planarians and their alleviation by sesquiterpene lactones, have the potential to contribute to a better understanding of how cocaine and related compounds cause their harmful effects in vertebrates and will likely point to potential strategies to prevent such undesirable effects in vivo.

**Figure 5.**
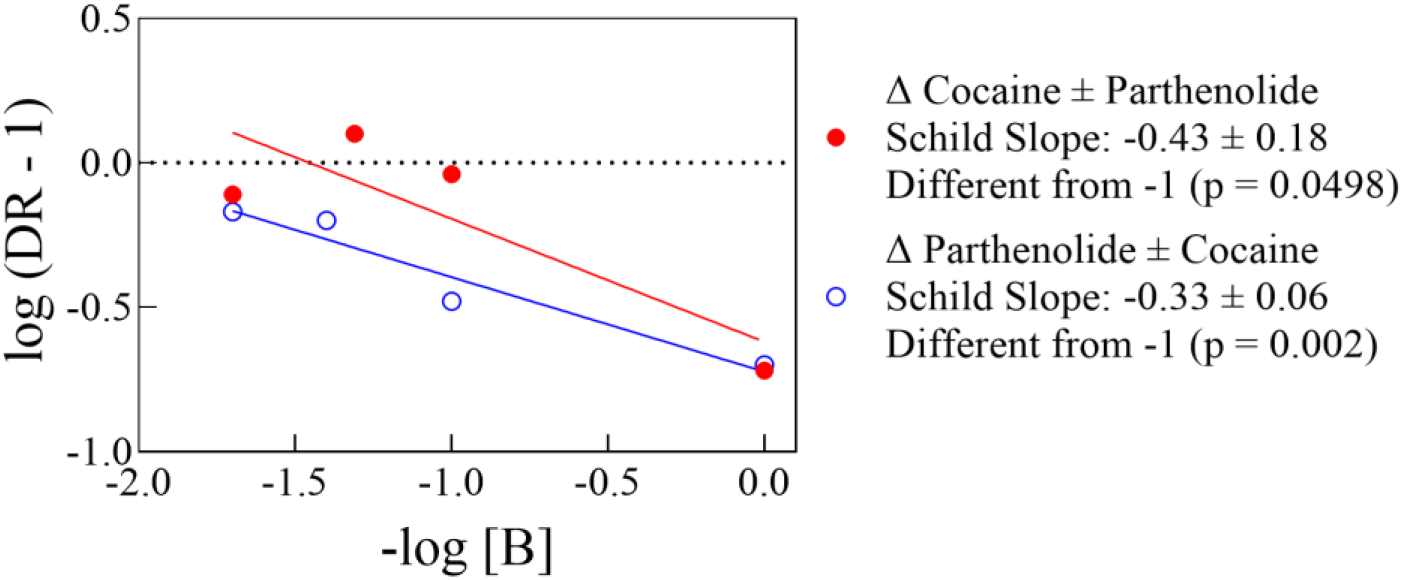
Schild Plots (abscissa and ordinate values from Table 1, see text). The Schild Slopes are not statistically different from each other (see text).

**Figure 6.**
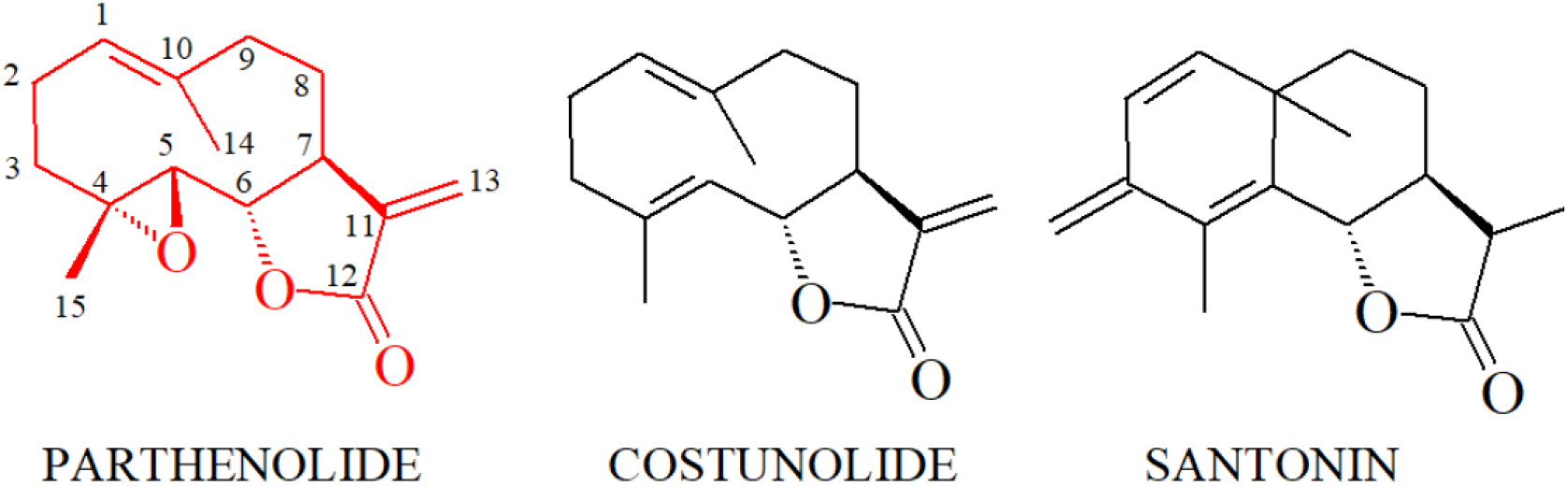
Structural comparison of parthenolide, costunolide, and santonin (see text).

## Author Contributions

Conceptualization, J.K., M.O., O.R.P.; methodology, J.K., M.O., O.R.P.; formal analysis, ORP; investigation, J.K., M.O., O.R.P.; resources, O.R.P.; data curation, O.R.P.; writing—original draft preparation, J.K; writing—review and editing, O.R.P.; supervision, O.R.P.; project administration, O.R.P.; funding acquisition, O.R.P. All authors read and agreed to the published version of the manuscript.

## Data Availability Statement

Available upon request.

## Acknowledgments

We thank the Department of Biology and the College of Science and Mathematics, West Chester University, for institutional and financial support.

